# DECREASING TRANSMURAL DISPERSION OF VENTRICULAR ACTION POTENTIAL REPOLARIZATION WITH MULTICHANNEL PHARMACOLOGY

**DOI:** 10.1101/2025.08.07.669124

**Authors:** Candido Cabo

## Abstract

Dispersion of repolarization results from a non-homogeneous recovery of excitability in cardiac tissue, and it is an important factor in arrhythmogenesis because it could lead to the initiation and maintenance of a variety of arrhythmias. Antiarrhythmic agents that prolong APD by selectively blocking specific ion channels (like I_Kr_) often increase dispersion of repolarization, which could result in a pro-arrhythmic risk. In this report, using computer models of the action potential of human epicardial and mid-myocardial myocytes, we have identified two strategies to prolong APD while reducing transmural dispersion of repolarization. The first strategy, which involves blocking several depolarizing and repolarizing ion channels (I_NaL_, I_CaL_, I_Kr_ and I_NaCa_), can reduce the transmural APD dispersion by about 20%. The second strategy, which involves the use of a combination of ion channel blockers and activators, results in a stronger reduction in transmural dispersion of repolarization than using only ion channel blockers. Enhancing I_Ks_ and blocking I_Kr_ can reduce transmural APD dispersion by about 70%. Our results suggest that a multichannel pharmacology strategy (as opposed to a single channel strategy), possibly using ion channel blockers and activators, can be effective at increasing APD while minimizing dispersion of repolarization.

## INTRODUCTION

Dispersion of repolarization results from a non-homogeneous recovery of excitability in cardiac tissue and can be influenced by various natural and pathological conditions (Surawicz 1989; Lukas 1997; Antzelevitch 2007, Antzelevitch 2008). Cells in the epicardial, mid-myocardial, and endocardial layers of the ventricles differ in their electrophysiological characteristics and their response to pharmacological agents (Lukas 1997; Antzelevitch 2008). In normal human hearts, dispersion of repolarization is generally small and considered benign (Kang et al 2017). However, there are genetic factors, acquired pathological conditions as well as unintended effects from antiarrhythmic drugs that may exacerbate the naturally occurring dispersion of repolarization (Shimizu and Antzelevitch 1998; Antzelevitch 2007; Antzelevitch 2008).

Dispersion of repolarization is an important factor in arrhythmogenesis, and it could lead to the initiation and maintenance of a variety of arrhythmias including Torsade de Pointes (TdP) and atrial fibrillation (AF) (Kuo et al 1985; Surawicz 1989; Antzelevitch 2005; Antzelevitch 2008; Avula et al 2019). Increased transmural dispersion of repolarization, frequently due to preferential prolongation of the APD of mid-myocardial cells, provides the arrhythmogenic substrate for TdP in patients with acquired or congenital LQT syndrome (Surawicz 1989; Antzelevitch 2005; Antzelevitch 2008). In that substrate, a premature ventricular contraction could lead to unidirectional block and initiation of reentrant waves (Belardinelli et al 2003; Antzelevitch 2005; Antzelevitch 2007). Human patients with paroxysmal or persistent AF show spatial heterogeneities in APD (Dicker et al 1998; Li et al. 2001; Avula et al. 2019), suggesting that dispersion of repolarization may provide a substrate for the initiation and maintenance of AF (Avula et al 2019).

Antiarrhythmic agents can have a proarrhythmic risk if they increase dispersion of repolarization. Class I antiarrhythmic drugs block the sodium channel but can also affect other ion channels, and they may increase or decrease dispersion of repolarization depending on their specific mechanism of action (Surawicz 1989; Belardinelli et al 2003). Class IA drugs, like quinidine and procainamide, block both sodium and potassium channels prolonging APD and are generally associated with an increased dispersion of repolarization and a higher risk of TdP (Surawicz 1989; Antzelevitch 2008), but not always (Milberg et al 2007). Class IB agent mexiletine blocks inactivated sodium channels and reduces both APD and dispersion of repolarization (Shimizu and Antzelevitch 1997). Class IC agent flecainide, similarly to some Class IA agents, tends to increase dispersion and contribute to arrhythmogenesis (Antzelevitch 2008). The impact of these drugs on dispersion of repolarization is a critical factor in their proarrhythmic risk (Antzelevitch 2005).

Class III antiarrhythmic drugs primarily act by blocking potassium channels, leading to APD prolongation to prevent reentrant arrhythmias (Peters et al 2020). Many Class III antiarrhythmic drugs, like sotalol and dofetilide, which act by blocking the rapid delayed rectifier ion channel (I_Kr_), tend to increase dispersion of repolarization by preferentially prolonging the APD of mid-myocardial cells (Lukas 1997; Antzelevitch 2008) and creating a substrate for reentrant arrhythmias like TdP (Antzelevitch 2005). However, there are class III agents that prolong APD homogeneously without an increase in dispersion of repolarization. For example, chromanol 293B, which blocks the slow delayed rectifier current (I_Ks_), prolongs APD homogenously without an increase in dispersion of repolarization (Antzelevitch 2005; Antzelevitch 2007). Amiodarone, a multi-channel blocker often classified as a class III antiarrhythmic agent, has been shown to increase APD while reducing dispersion of repolarization by prolonging APD in endocardial and epicardial cells but not in mid-myocardial cells (Sicouri et al 1997; Drouin et al 1998; Vasallo and Trohman 2007; Arpadffy et al 2020; Gelman et al 2024).

Other heart acting drugs like anti-anginal and adrenergic agents may also cause a decrease in dispersion of repolarization. Anti-anginal agent ranolazine, which acts on sodium, calcium and potassium channels, has a similar electrophysiological effect to amiodarone by prolonging APD in epicardial cells but not in mid-myocardial cells, and therefore reducing dispersion of repolarization (Antzelevitch et al 2004; Hasenfuss and Maier 2008). Carvedilol is an alpha- and beta-adrenergic antagonist that also modulates potassium, sodium and calcium channels (Karle et al 2001). In a rabbit model of congestive heart failure, carvedilol causes a reduction of dispersion of repolarization by prolonging APD in epicardial and endocardial cells to a larger extent than in mid-myocardial cells (Zhong et al 2007). Anesthetics like sodium pentobarbital and propofol have also been shown to reduce dispersion of repolarization by prolonging APD in epicardial and endocardial cells but to a lesser extent in mid-myocardial cells (Shimizu et al 1999; Ellerman et al 2020).

In summary, while APD prolongation often results in an increased dispersion of repolarization, it is the increase in dispersion itself that is considered the primary arrhythmogenic substrate for TdP (Antzelevitch 2005; Antzelevitch 2008; Belardinelli et al 2003; Arpadffy et al. 2020). Pharmacological agents that decrease dispersion of repolarization (like amiodarone, ranolazine and pentobarbital) often achieve this through the modulation of multiple ion channels, by having a differential effect in different myocardial cell types resulting in a more homogeneous recovery of excitability. This reduction in dispersion is considered crucial for mitigating the risk of serious arrhythmias (Antzelevitch et 2004; Antzelevitch 2005; Antzelevitch 2007; Antzelevitch 2008; Trenor et al 2013; Arpadffy et al 2020; Ellerman et al. 2020). In this report, we investigate, using a computer model of human ventricle, the mechanisms by which multi-channel pharmacology can reduce transmural dispersion of repolarization.

## METHODS

### Computer models of the action potential

We simulated the cardiac action potential using the ORd (O’Hara et al. 2011) models of human ventricular epicardial and mid-myocardial cells. Both models are publicly available from the Rudy Lab web site (https://rudylab.wustl.edu/code-downloads/).

We investigated changes in transmural heterogeneities between the epicardial and mid-myocardial action potential models using ion channel blockers and enhancers by modulating the maximum conductance of: late sodium current (I_NaL_; range: 0-2x control), L-type calcium current (I_CaL_; range: 0.5-1.5x control), slow delayed rectifier potassium current (I_Ks_; range: 0-50x control), rapid delayed rectifier potassium current (I_Kr_; range: 0-2x control), inward rectifier potassium current (I_K1_; range: 0.2-2x control), sodium/potassium pump (I_NaK_; range: 0.5-1.5x control) and sodium/calcium exchanger (I_NaCa_; range: 0.5-1.5x control). For the simulations using only ion channel blockers we limited the maximum conductance to that of control. Action potentials were initiated with a depolarizing current with a strength 1.5x the stimulation threshold. We report measurements on action potentials that were calculated after 30 minutes of stimulation to achieve steady-state (Cabo 2022).

### Action potential features

The phases of the action potential were quantified as described in an earlier report (Cabo 2022). In short, phase 1 begins at the time of action potential depolarization and it ends at the time repolarization starts, which is when the total ion current becomes positive (Cabo 2022, Figure 1). Phase 2 starts when phase 1 ends, and it ends when I_K1_ rises to 10% of its peak (Cabo 2022). In the ORd model the end of phase 2 occurs when the membrane repolarizes to -39 mV. Phase 3 starts at the end of phase 2, and it ends when the action potential repolarizes by 90% of the action potential amplitude from its maximum depolarization potential.

**Figure 1.**
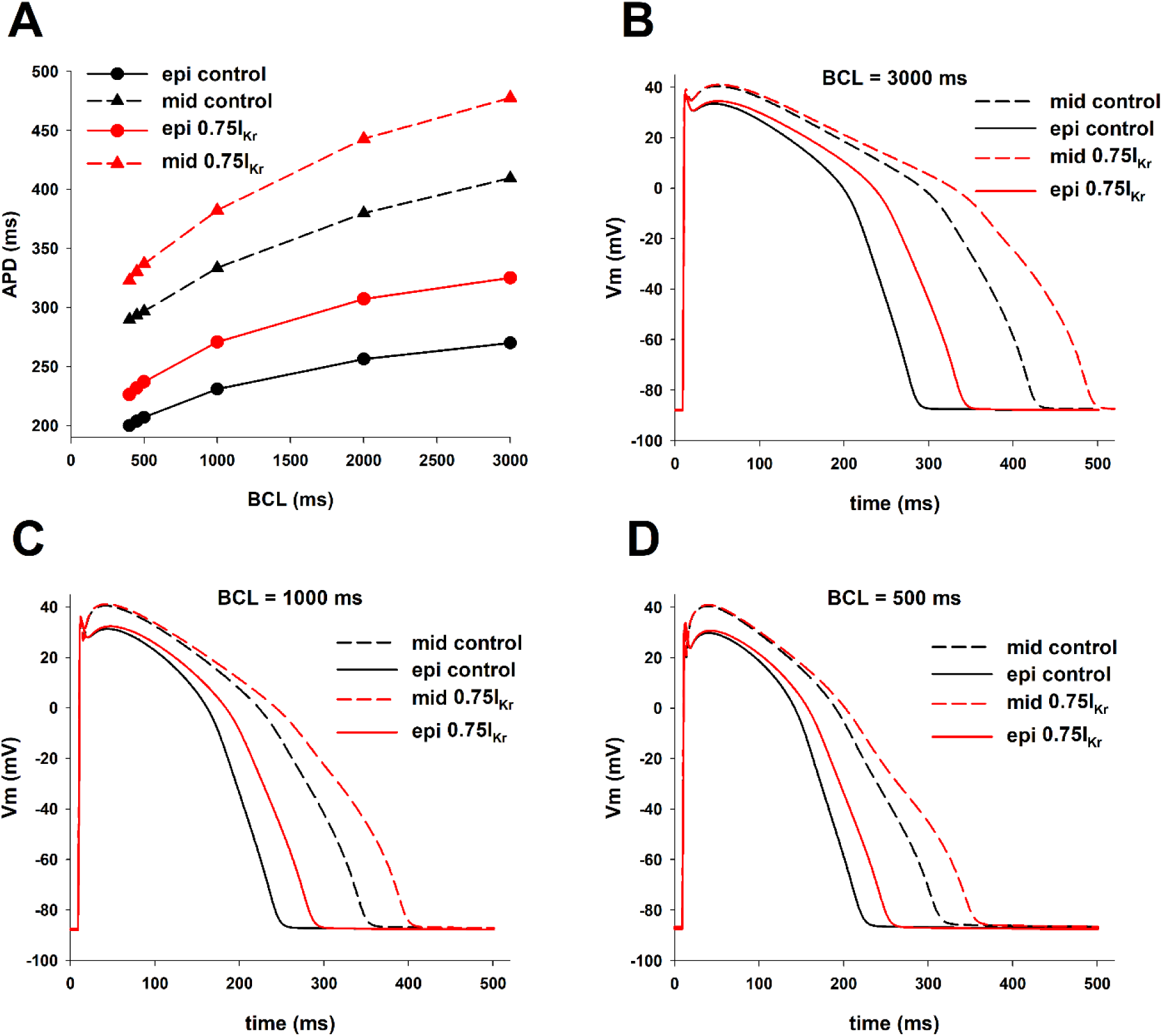
Dispersion of repolarization between epicardial and mid-myocardial cells. **A:** Action potential duration (APD) during control (black) and 25% block of I_Kr_ (red), for epicardial (circles, solid lines) and mid-myocardial cells (triangles, dashed lines), for different BCLs. **B:** Action potentials for BCL = 3000 ms. **C:** Action potentials for BCL = 1000 ms. **D:** Action potentials for BCL = 500 ms. See text for detailed description.

### Particle Swarm Optimization algorithm

As before (Cabo 2022), we used the Particle Swarm Optimization (PSO) algorithm (Kennedy and Eberhart, 1995) to find the optimal combination of maximum conductance of I_NaL_, I_CaL_, I_Ks_, I_Kr_, I_K1_ I_NaK_, and I_NaCa_ to minimize the difference between APD in epicardial and mid-myocardial cells. We used an implementation of the PSO algorithm publicly available in the Github repository (https://github.com/kkentzo/pso). The goal of the PSO optimization algorithm was to minimize the APD differences between epicardial and mid-myocardial cells, and differences between the APD of either epicardial or mid-myocardial cells and a given APD target set by the user, depending on the optimization simulation. The ion currents and the range of variation (minimum and maximum values) of ion channel maximum conductance allowed for the optimization process are specified above in section *Computer Models of the Action Potential*.

### Backward Feature Elimination

We used a backward feature elimination procedure to investigate the relative contribution of each ion current to the transmural heterogeneities (dispersion) in the action potential (Cabo 2023). After applying the PSO optimization to find the combination of maximum conductance of I_NaL_, I_CaL_, I_Ks_, I_Kr_, I_K1_, I_NaK_, and I_NaCa_ that minimize APD dispersion, optimization was applied to the seven possible subsets of six currents (i.e., [I_CaL_, I_Ks_, I_Kr_, I_K1_, I_NaK_, and I_NaCa_], [I_NaL_, I_Ks_, I_Kr_, I_K1_, I_NaK_, and I_NaCa_], [I_NaL_, I_Ca_, I_Kr_, I_K1_, I_NaK_, and I_NaCa_], [I_NaL_, I_CaL_, I_Ks_, I_K1_, I_NaK_, and I_NaCa_], [I_NaL_, I_CaL_, I_Ks_, I_Kr_, I_NaK_, and I_NaCa_], [I_NaL_, I_CaL_, I_Ks_, I_Kr_, I_K1_, and I_NaCa_], [I_NaL_, I_CaL_, I_Ks_, I_Kr_, I_K1_, and I_NaK_]). The current not present in each subset was kept at the control value. The subset resulting in the larger reduction of dispersion after PSO optimization was selected for the next step in the elimination procedure. The ion current not present in the selected subset was the current that contributed less to a reduction in APD dispersion, and it was consequently eliminated (i.e., its maximum conductance was set to the control value). This process of elimination was repeated until only two ion currents were left.

## RESULTS

### Transmural action potential heterogeneity

Figure 1A shows the differences in APD between epicardial (solid black circles; solid black line) and mid-myocardial (solid black triangles; dashed black line) cells, for BCLs between 400 ms and 3000 ms under physiological conditions (control). For each BCL, APD of mid-myocardial cells is larger than APD in epicardial cells. Differences in APD (transmural APD dispersion) range from 139 ms at BCL = 3000 ms to 90 ms at BCL = 400 ms (Figure 1A). Figure 1A also shows that selective block of I_Kr_ increases APD dispersion by increasing APD in mid-myocardial cells to a larger extent than APD in epicardial cells (epicardial cells: red circles, solid red line; mid-myocardial cells: red triangles, dashed red line). With 25% I_Kr_ block, APD dispersion increased from 139 ms to 152 ms at BCL = 3000 ms (∼9% increase), and from 90 ms to 97 ms at BCL = 400 ms (∼8% increase). Action potentials for epicardial and mid-myocardial cells during control and 25% I_Kr_ block for three BCLs are shown in Figures1B-1D.

Figure 2 shows the major depolarizing and repolarizing ion currents during the action potential for epicardial (black) and mid-myocardial (red) cells during stimulation with BCL = 1000 ms during control. Channel density of depolarizing currents, I_CaL_ and I_NaL,_ in mid-myocardial cells is about twice that of epicardial cells (O’Hara et al. 2011), which explain the larger I_CaL_ and I_NaL_ currents in mid-myocardial cells and the more positive depolarization in the action potential of mid-myocardial cells during phases 1 and 2 of the action potential (Figure 2, top left). The Na/Ca exchanger (I_NaCa_) depolarizing current during phase 2 and phase 3 repolarization is larger for mid-myocardial than for epicardial cells, consistent with the 30% larger channel density of the exchanger in mid-myocardial cells (O’Hara et al. 2011). Therefore, I_CaL_, I_NaL_ and I_NaCa_ contribute to the more positive depolarization and longer APD in mid-myocardial than in epicardial cells. The channel density of I_Kr_ in epicardial cells is about 50% larger than in mid-myocardial cells (O’Hara et al. 2011), which results in the much larger repolarizing current in epicardial cells (Figure 2, top right). Like for I_Kr_, the channel density of I_Ks_ in epicardial cells is also about 50% larger than in mid-myocardial cells (O’Hara et al. 2011). However, in contrast to what happened with I_Kr_, the repolarizing I_Ks_ current is larger for mid-myocardial cells than for epicardial cells despite its smaller channel density (Figure 2, right, second plot from the top). I_Ks_ activates more slowly and at more positive transmembrane potentials than I_Kr_ (Liu and Antzelevitch 1995). Since mid-myocardial cells depolarize to more positive potentials and stay depolarized longer at positive potentials than epicardial cells, I_Ks_ is larger in mid-myocardial cells (Figure 2). There are essentially no differences in I_K1_ between epicardial and mid-myocardial cells (Figure 2, right, third plot from the top). Channel density of I_NaK_ is smaller in mid-myocardial than epicardial cells (O’Hara et al. 2011) but I_NaK_ current during phase 2 and phase 3 repolarization is larger in mid-myocardial cells than in epicardial cells (Figure 2, right bottom), as a result of an increase in [Na]_i_ (from 7.8mM to 8.9mM) (Glitsch 2001). Still, the contribution of I_Ks_ and I_NaK_ to repolarization is much smaller than that of I_Kr_ (Figure 2), and overall, the total repolarizing current is much larger in epicardial than in mid-myocardial cells, which results in the shorter APD. In summary, the shorter APD in epicardial cells is mainly due to their larger I_Kr_, which results in a stronger total repolarizing current.

**Figure 2.**
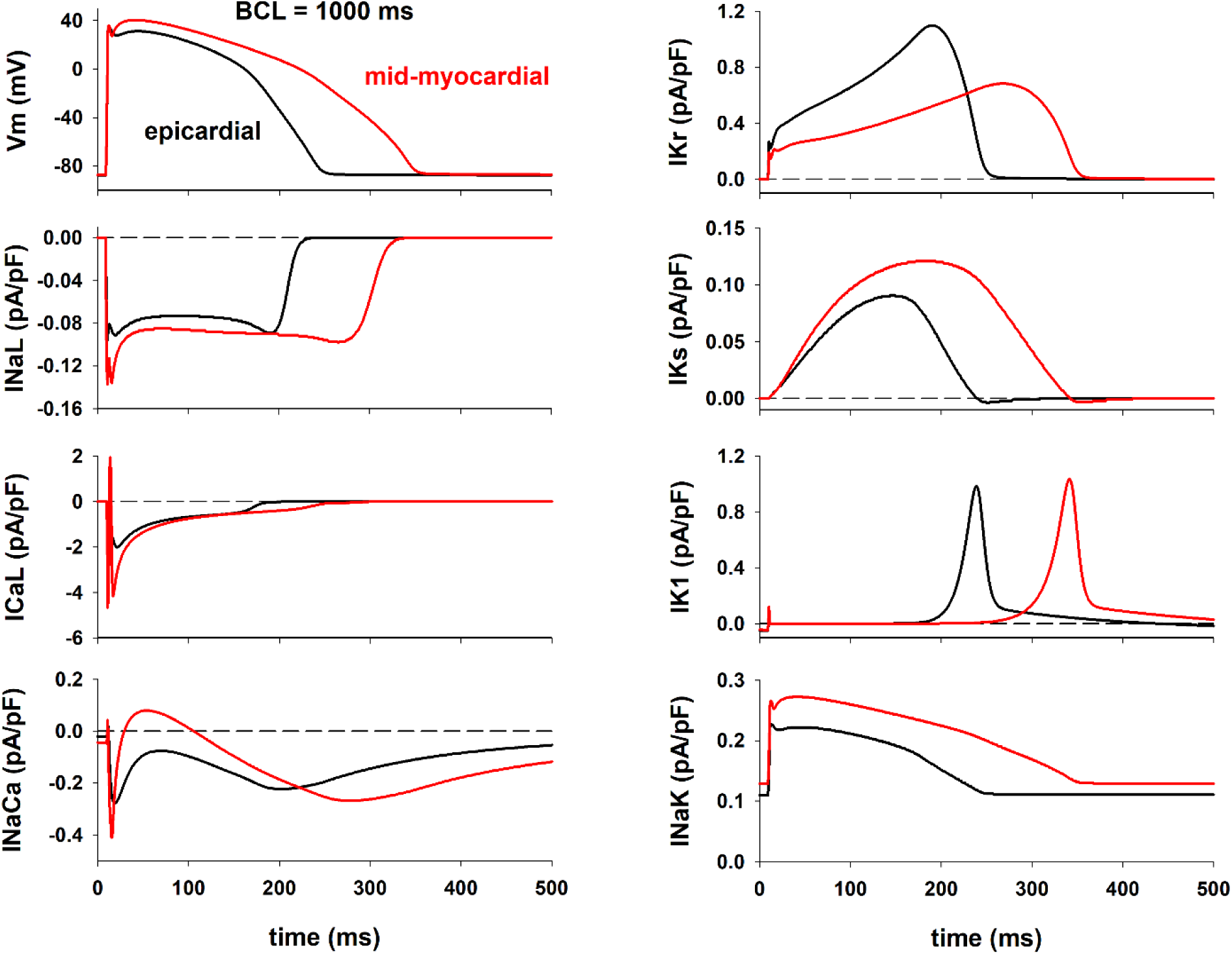
Action potentials (top left) as well as depolarizing and repolarizing ion currents during the action potential for epicardial (black) and mid-myocardial (red) cells during stimulation with BCL = 1000 ms during control. See text for detailed description.

### Rate dependence of transmural heterogeneity

Figure 1 shows that transmural heterogeneities in APD between mid-myocardial and epicardial cells are rate dependent: a decrease in BCL leads to a decrease in APD dispersion (Figure 1A, black solid and dashed lines). Figure 3A shows that for both epicardial and mid-myocardial cells, the decrease in APD with BCL is caused by a decrease in the duration of phase 2&3 (i.e. the total duration of phase 2 and phase 3) repolarization. Figure 3B shows the relationship between average total ion current during phase 2&3 repolarization of the action potential (I_tot_) and the duration of phase 2&3 repolarization for epicardial and mid-myocardial cells. In both cells, such a relationship can be modeled by a hyperbola, I_ion,ph2ph3_ = K / (duration of phase 2&3 repolarization), where K is the approximate change in transmembrane potential during phase 2&3 repolarization (Cabo 2023). For both, epicardial and mid-myocardial cells, a decrease in BCL from 3000 ms to 500 ms results in an increase in average I_tot_ (27% epi; 38% mid), I_dep_ (23% epi; 24% mid) and I_rep_ (25% epi; 29% mid) (black and white bars in Figure 3C and 3D). The increase in depolarizing current is a consequence of an increase in I_CaL_ and I_NaCa_ for both types of cells (I_CaL_ and I_NaCa_ in Figures 3C and 3D). The increase in repolarizing current is a consequence of the increase in I_NaK_ (I_NaK_ in Figures 3C and 3D). The increase in average I_tot,ph2ph3_ when BCL is reduced from 3000 ms to 500 ms is larger for epicardial (0.14 pA/pF) than for mid-myocardial (0.12 pA/pF) cells (Figures 3B, 3C and 4D), but that increase results in a larger reduction in the duration of phase 2&3 for mid-myocardial (102 ms) than for epicardial cells (57 ms). This is a consequence of the hyperbolic relationship between average total ion current during repolarization (I_tot,ph2ph3_) and the duration of phase 2&3 (solid line, Figure 3B); the hyperbola has a larger slope for longer values of phase 2&3 (mid-myocardial cells), than for shorter values of phase 2&3 (epicardial cells).

**Figure 3.**
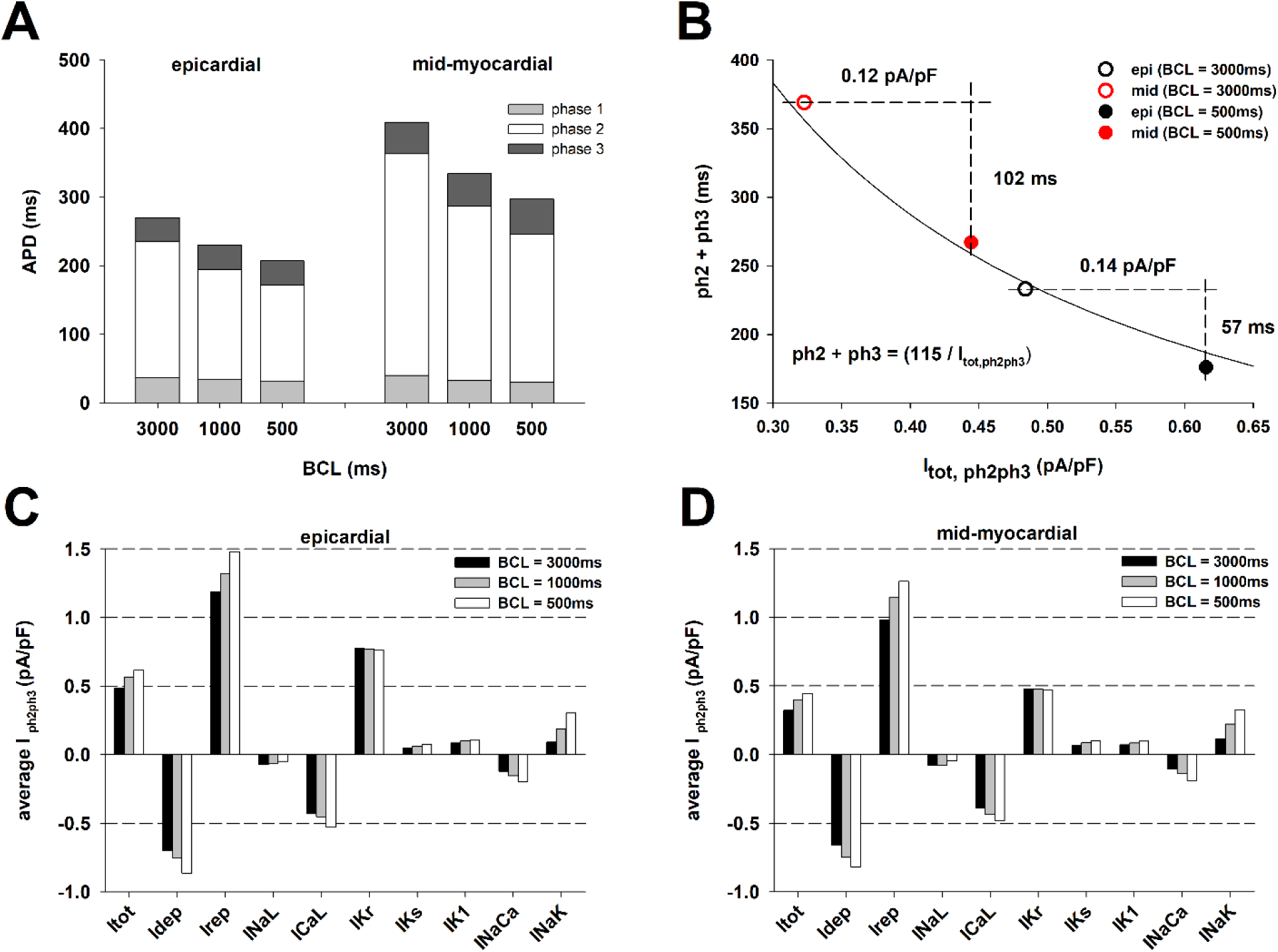
Rate dependance of dispersion of repolarization between epicardial and mid-myocardial cells. A: Duration of phase 1, 2 and 3 of the action potential for epicardial and mid-myocardial cells for different BCLs. B: Relationship between the duration of phase 2 and 3 and the average total ion current during phase 2 and 3. C: Average ion currents during phase 2 and 3 of the action potential for epicardial cells for different BCLs. D: Average ion currents during phase 2 and 3 of the action potential for mid-myocardial cells for different BCLs. See text for detailed description.

### Reduction of transmural action potential heterogeneity with ion channel blockers

We used an optimization algorithm to investigate the optimal combinations of ion channel blockers that result in a reduction of APD dispersion. Figure 4 shows the results of an intervention that could reduce APD dispersion by increasing APD in epicardial cells while keeping APD in mid-myocardial cells close to the control value by blocking several depolarizing and repolarizing currents (I_NaL_, I_CaL_, I_Kr_, I_NaK_ and I_NaCa_). The intervention caused a reduction of APD dispersion from 139 ms to 112 ms (∼19% reduction) at BCL = 3000 ms, and from 90 ms to 73 ms (∼19% reduction) at BCL = 400 ms (Figure 4A). The decrease in APD dispersion with optimal multichannel block resulted from a preferential increase in APD of epicardial cells. Figures 4B- 4D show the corresponding action potentials at selected BCLs. In contrast, the selective 25% block of I_Kr_, which achieves a similar prolongation of APD in epicardial cells than the intervention in Figure 4, caused an 8-9% *increase* of APD dispersion (Figure 1A). The increase in APD dispersion with the selective I_Kr_ block resulted from a preferential prolongation of APD of mid-myocardial cells. The optimal multichannel block shown in Figure 4 decreases APD dispersion by a preferential prolongation of APD of epicardial cells.

**Figure 4.**
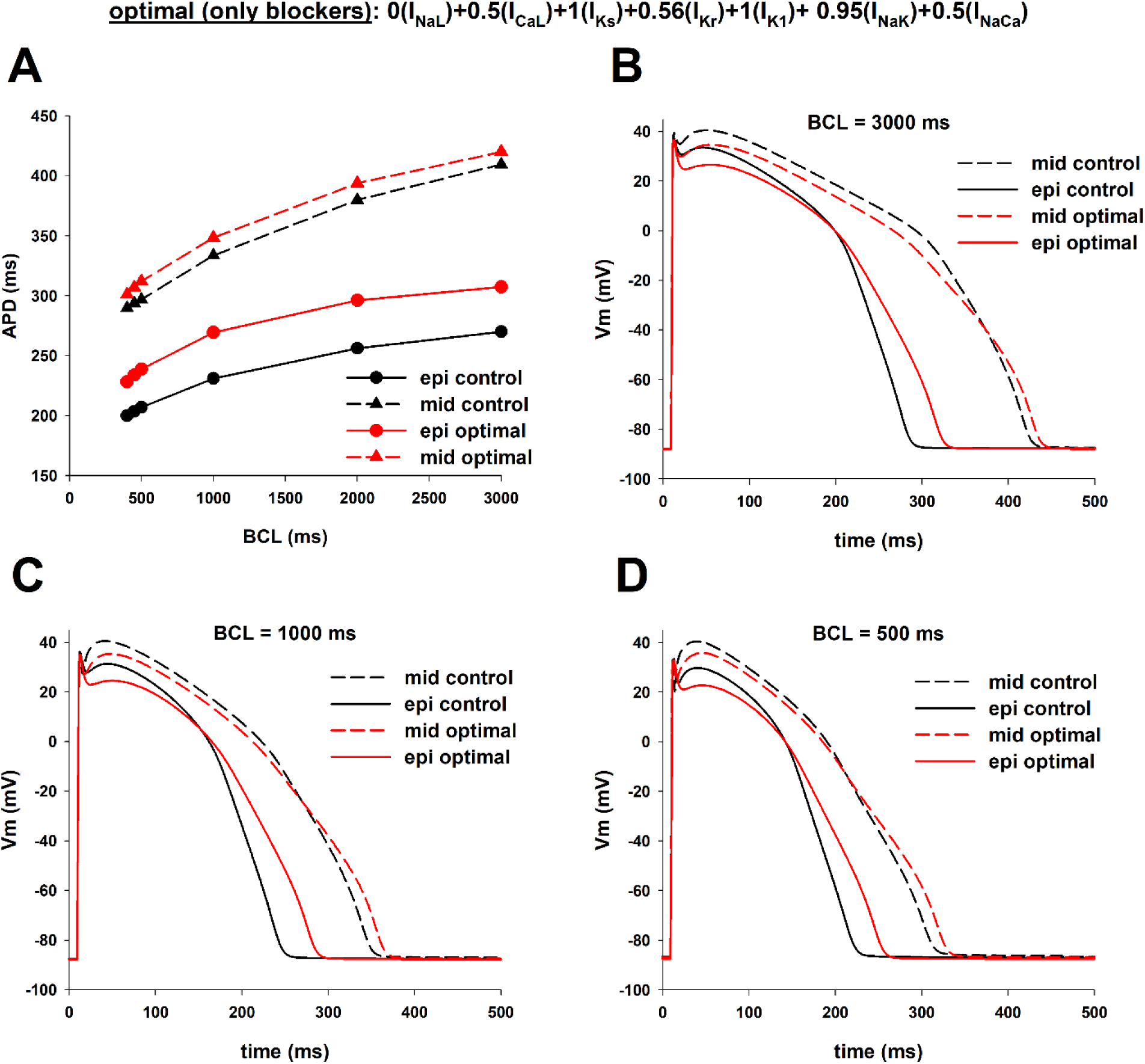
Optimal combination of ion channel blockers that results in a reduction of dispersion of repolarization while keeping APD of mid-myocardial cells close to control. **A:** The intervention reduced APD dispersion by increasing APD in epicardial cells while keeping APD in mid-myocardial cells close to the control value by blocking several depolarizing and repolarizing currents (I_NaL_, I_CaL_, I_Kr_, I_NaK_ and I_NaCa_). Panels **B-D** show the corresponding action potentials of different BCLs. The format of the figure is the same as Figure 1. See text for detailed description.

Figure 5 shows action potential features and average currents during phase 2&3 repolarization for control, selective 25% block of I_Kr_, and optimal multichannel block when BCL = 1000 ms. The effect of both interventions on the duration of phase 1 is much smaller than their effect on the duration of phase 2 and phase 3 for both types of cells (Figure 5A). Selective 25% block of I_Kr_ causes a reduction of average I_tot_ of 0.09 pA/pF in epicardial cells and of 0.05 pA/pF in mid-myocardial cells (Figures 5B, 5C and 5D). The block of depolarizing currents (I_NaL_, I_CaL_, and I_NaCa_) causes about the same reduction of average depolarizing current in epicardial cells (0.11 pA/pF) and mid-myocardial cells (0.09 pA/pF) (I_dep_ in Figures 5C and 5D). The larger reduction of average I_tot_ in epicardial cells is a consequence of the larger reduction of I_Kr_ in epicardial cells (0.18 pA/pF, Figures 5B and 5C) than in mid-myocardial cells (0.11 pA/pF, Figures 5B and 5D). However, despite the larger reduction of I_tot_ in epicardial cells than in mid-myocardial cells, the increase in duration of phase 2&3 repolarization (and APD) is smaller in epicardial than in mid-myocardial cells. This is a consequence of the hyperbolic relationship between average I_tot_ current during phase 2&3 repolarization (I_tot,ph2ph3_) and the duration of phase 2&3 repolarization; the hyperbola has a larger slope for action potentials with longer phase 2&3 (mid-myocardial cells), than for action potentials with shorter phase 2&3 (epicardial cells).

**Figure 5.**
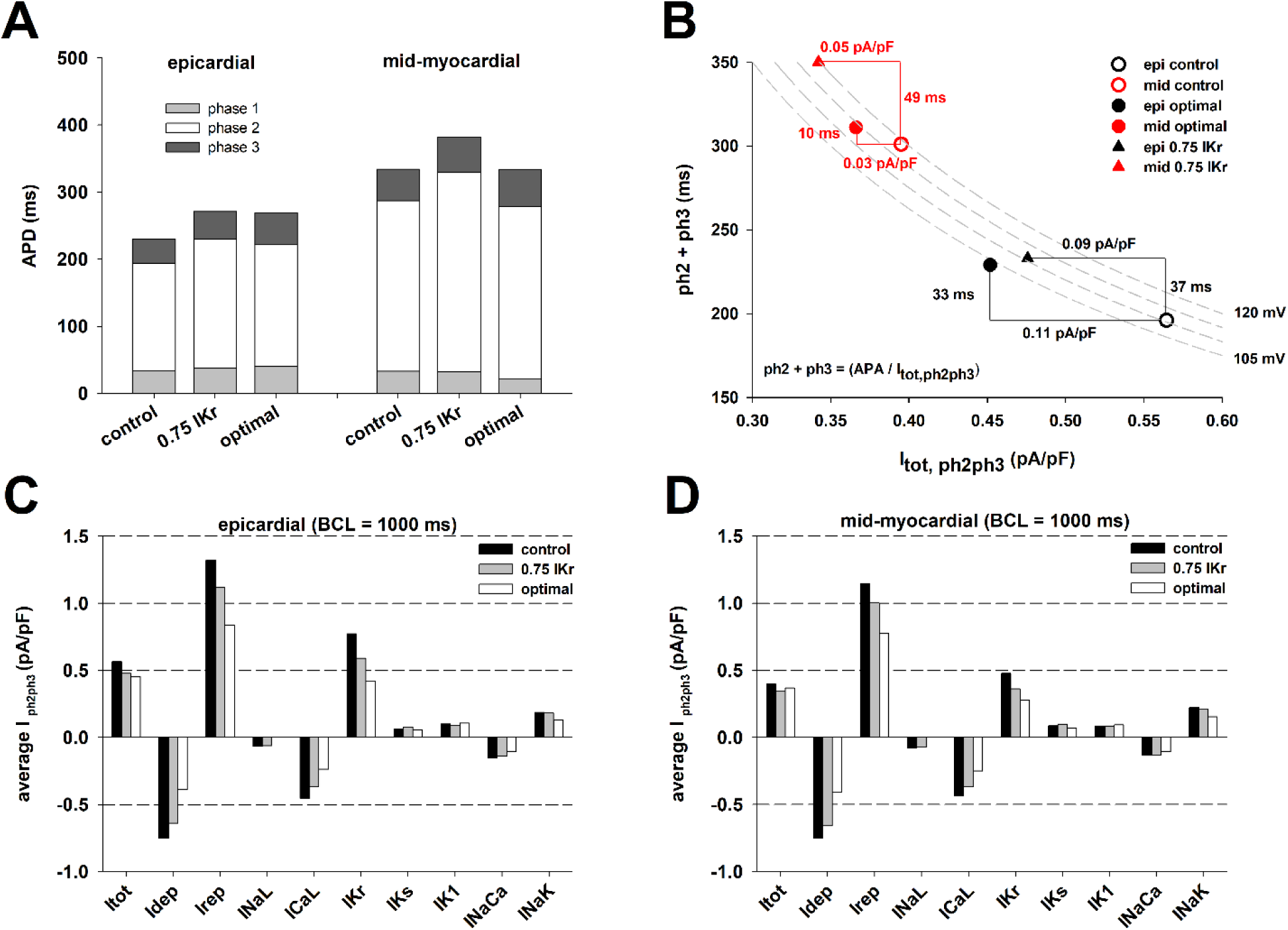
Comparison of action potential features for epicardial and mid-myocardial cells during control, 25% block of IKr, and the optimal multichannel block (optimal) in Figure 4 that causes a reduction in dispersion of repolarization with BCL = 1000 ms. A: Duration of phase 1, 2 and 3 for epicardial and mid-myocardial cells. B: Relationship between the duration of phase 2 and 3 and the average total ion current during phase 2 and 3 during control, 25% block of IKr, and the optimal intervention. C: Average ion currents during phase 2 and 3 of the action potential for epicardial cells during control, IKr block and the optimal intervention. D: Average ion currents during phase 2 and 3 of the action potential for mid-myocardial cells during control, I_Kr_ block and the optimal intervention. See text for detailed description.

Optimal block of several depolarizing and repolarizing ion channels increases APD in epicardial cells by 38 ms (from 231 ms to 269 ms) and APD of mid-myocardial by 14 ms (from 334 ms to 348 ms), which results in a decrease in transmural APD dispersion from 103 ms to 79 ms (∼23% reduction) (Figure 4C and Figure 5A). Optimal multichannel block causes a larger reduction in average I_tot_ in epicardial cells (0.11 pA/pF) than in mid-myocardial cells (0.03 pA/pF), resulting in the larger increase in the duration of phase 2&3 repolarization (and APD) in epicardial cells (Figures 5B, 5C and 5D). The block of depolarizing currents (I_NaL_, I_CaL_, and I_NaCa_) causes about the same reduction of average depolarizing current in epicardial cells (0.37 pA/pF) and in mid-myocardial cells (0.34 pA/pF) (I_dep_ in Figures 5C and 5D). However, there is a larger decrease in average repolarizing currents in epicardial cells (0.48 pA/pF) than in mid-myocardial cells (0.37 pA/pF) (I_rep_ in Figures 5C and 5D). That is a consequence of the larger reduction of I_Kr_ in epicardial cells (0.35 pA/pF, Figures 5B and 5C) than in mid-myocardial cells (0.20 pA/pF, Figures 5B and 5D) and the larger contribution of I_Kr_ to the total repolarizing current in epicardial cells than in mid-myocardial cells (I_rep_ in Figures 5C and 5D).

Figure 6 shows the results of an intervention that decreases APD dispersion by reducing APD in mid-myocardial cells while keeping APD in epicardial cells close to the control value. The intervention caused a reduction of APD dispersion from 139 ms at BCL = 3000 ms to 104 ms (∼25% reduction), and from 90 ms at BCL = 400 ms to 68 ms (∼24% reduction) (Figure 6A). Figures 6B-6D show the corresponding action potentials at selected BCLs. As with the intervention in Figure 4, the results suggest that the combined block of I_NaL_, I_CaL_, I_Kr_ and I_NaCa_ can lead to a substantial reduction in transmural APD dispersion.

**Figure 6.**
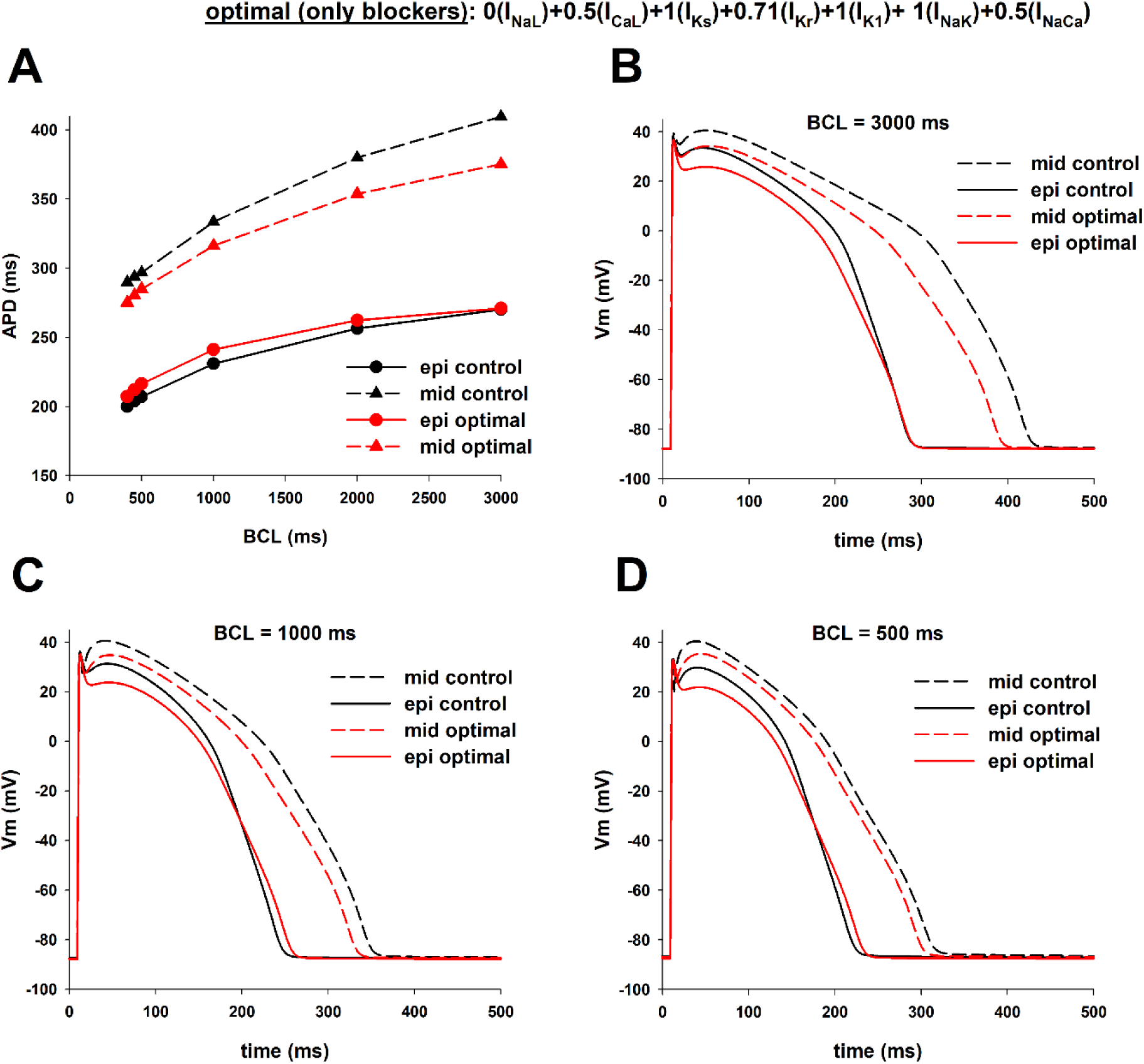
Optimal combination of ion channel blockers that results in a reduction of dispersion of repolarization while keeping APD of epicardial cells close to control. **A:** The intervention reduced APD dispersion by decreasing APD in mid-myocardial cells while keeping APD in epicardial cells close to the control value by blocking several depolarizing and repolarizing currents (I_NaL_, I_CaL_, I_Kr_ and I_NaCa_). Panels **B-D** show the corresponding action potentials of different BCLs. The format of the figure is the same as Figure 1. See text for detailed description.

To investigate the relative contribution of each ion current to the reduction of transmural APD dispersion shown in Figures 4 and 6, we used a backward feature elimination procedure (Figure 7). Figure 7A shows the results of the elimination procedure in the optimal multichannel block in Figure 4 on APD dispersion with BCL = 1000 ms. Optimal block of I_NaL_, I_CaL_, I_Kr_ and I_NaCa_ reduces APD dispersion from 103 ms (Figure 7A, control) to 75 ms (Figure 7A, step 1). Elimination of I_CaL_ block (i.e. using the same I_CaL_ as control) increases APD dispersion to 85 ms (Figure 7A, step 2). Further elimination of I_NaCa_ block increases APD dispersion to 95 ms (Figure 7A, step 3). The results in Figure 7A indicate that I_Kr_ and I_NaL_ are the two most important ion channels to reduce APD dispersion, but it also shows that block of I_CaL_ and I_NaCa_ contribute significantly to reduction of APD dispersion. Figure 7B shows the results of the feature elimination procedure in the optimal multichannel block in Figure 6 on APD dispersion with BCL = 1000 ms. Even though 5% block of I_NaK_ is part of the optimal combination of ion channels blockers to reduce APD dispersion (Figure 7B, step 1), its elimination does not significantly change the resulting APD dispersion (Figure 7B, step 2). Subsequent steps 2 to 4 in Figure 7B follow the same pattern as in Figure 7A, highlighting the importance of I_Kr_ and I_NaL_ in the reduction of APD dispersion, but also the important contribution of I_CaL_ and I_NaCa_.

**Figure 7.**
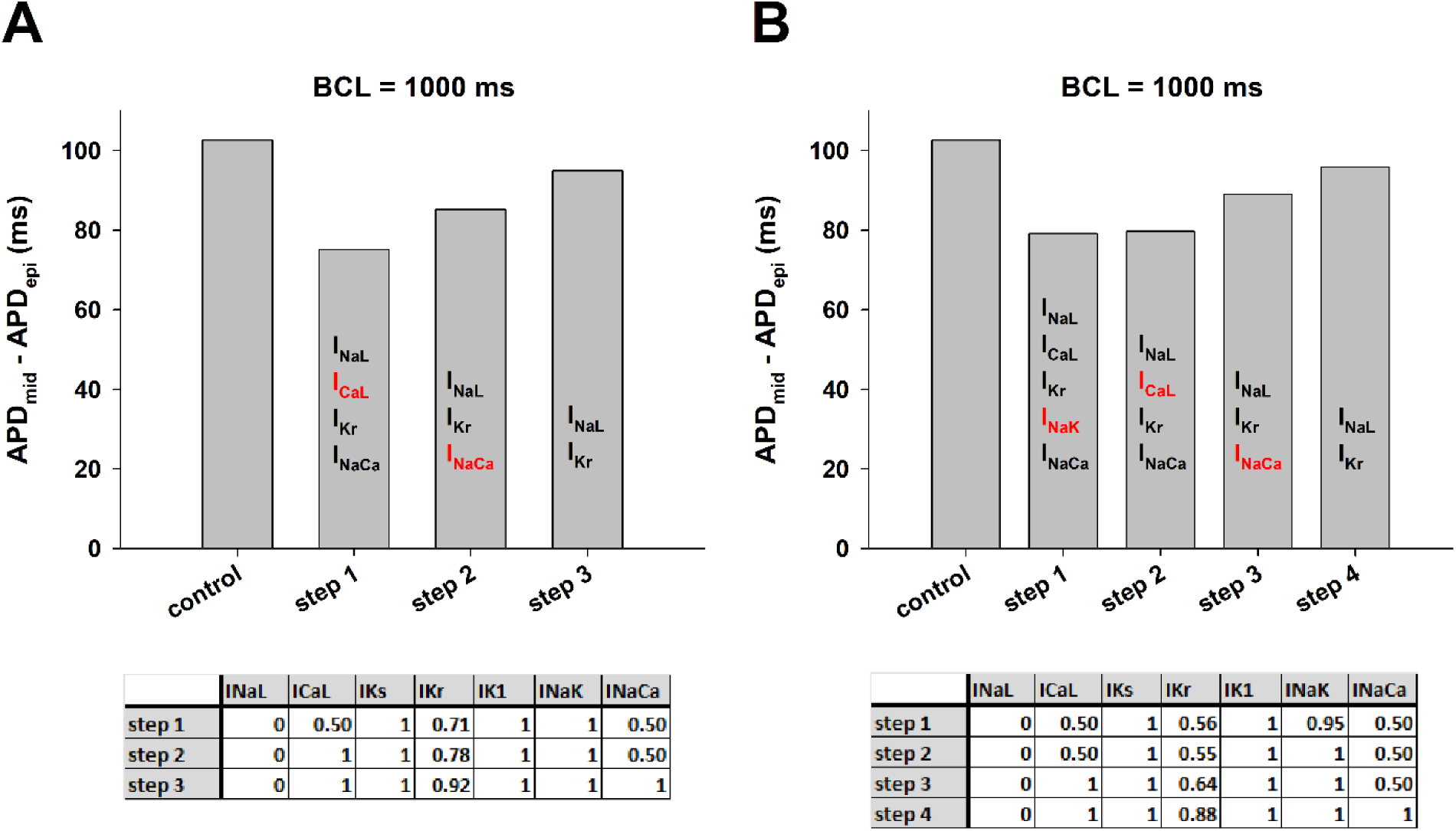
Backward feature elimination procedure applied to interventions in Figures 4 (panel A) and 6 (panel B) that reduce dispersion of repolarization when BCL = 1000 ms. The top of each panel shows APD dispersion (i.e. the difference between APD in epicardial and mid-myocardial cells) during control and after each step in the procedure. Each vertical bar shows the ion currents that were subject to block. The ion current in red indicates the current eliminated after that step in the procedure because it contributes less to reducing APD dispersion. The bottom of each panel shows the contribution of each ion current at a specific step. See text for detailed description.

### Reduction of transmural action potential heterogeneity with ion channel blockers and activators

We used an optimization algorithm to investigate the optimal combinations of ion channel blockers and activators that result in a reduction of APD dispersion. Figure 8 shows the results of an intervention that could reduce APD dispersion by reducing APD in mid-myocardial cells while keeping APD in epicardial cells close to the control value. The intervention caused a reduction of APD dispersion from 139 ms at BCL = 3000 ms to 27 ms (∼81% reduction), and from 90 ms at BCL = 400 ms to 16 ms (∼82% reduction) (Figure 8A). Figures 8B-8D show the corresponding action potentials at selected BCLs. In both epicardial and mid-myocardial cells enhancement of I_Ks_ and block of I_Kr_ leads to an acceleration of phase 2 repolarization and a deceleration of phase 3 repolarization (Figures 8B-8D). The reduction of transmural APD dispersion obtained by using an optimal combination of ion channel blockers and activators (Figure 8) is four times larger than the reduction obtained by using just ion channel blockers (Figures 4 and 6).

**Figure 8.**
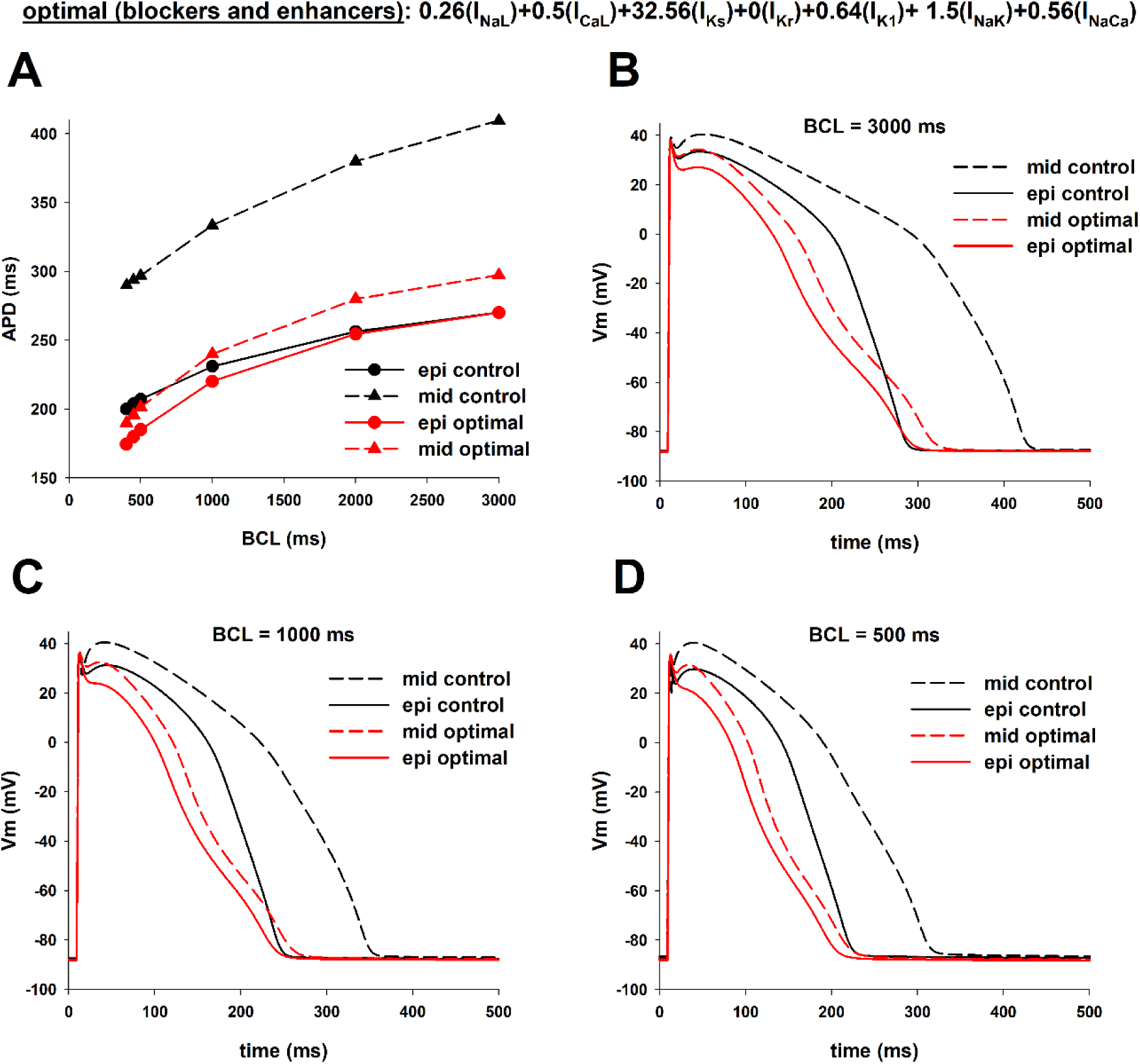
Optimal combination of ion channel blockers and activators that results in a reduction of dispersion of repolarization while keeping APD of epicardial cells close to control. **A:** The intervention reduced APD dispersion by decreasing APD in mid-myocardial cells while keeping APD in epicardial cells close to the control value by blocking and activating several depolarizing and repolarizing currents. Panels **B-D** show the corresponding action potentials of different BCLs. The format of the figure is the same as Figure 1. See text for detailed description.

To investigate the relative contribution of each ion current to the reduction of transmural heterogeneities in the action potential we used a backward feature elimination procedure (Figure 9). Transmural dispersion during control, with a BCL = 1000 ms, was 103 ms (Figure 9, control). The optimal combination of ion channel conductance of I_NaL_, I_CaL_, I_Ks_, I_Kr_, I_K1_, and I_NaK_ and I_NaCa_ reduces APD dispersion to 20 ms (Figure 8A, 8C and Figure 9, step 1). Not all ion channels contribute equally to the reduction of APD dispersion. Elimination of the modulation of I_K1_ and I_NaL_ (that is, keeping those currents the same as control) does not change the value of APD dispersion indicating that those ion channels do not contribute much to the reduction in APD dispersion (Figure 9, steps 2 and 3). Elimination of I_CaL_ increases APD dispersion modestly to 23 ms (Figure 9, step 4). Further elimination of I_NaL_ increases APD dispersion to 25 ms (Figure 9, step 5). Additional elimination of I_NaCa_ increases APD dispersion to 34 ms (Figure 9, step 6). These results show the importance of enhancing I_Ks_ and blocking I_Kr_ to obtain a strong reduction in APD dispersion; additional modulation of I_NaCa_, I_NaL_ and I_CaL_ can further decrease APD dispersion. All in all, the results in Figure 9 show that the currents that contribute the most to APD dispersion reduction are I_Ks_ and I_Kr_; modulation of just those two currents reduce APD dispersion from 103 ms to 34 ms at BCL = 1000ms.

**Figure 9.**
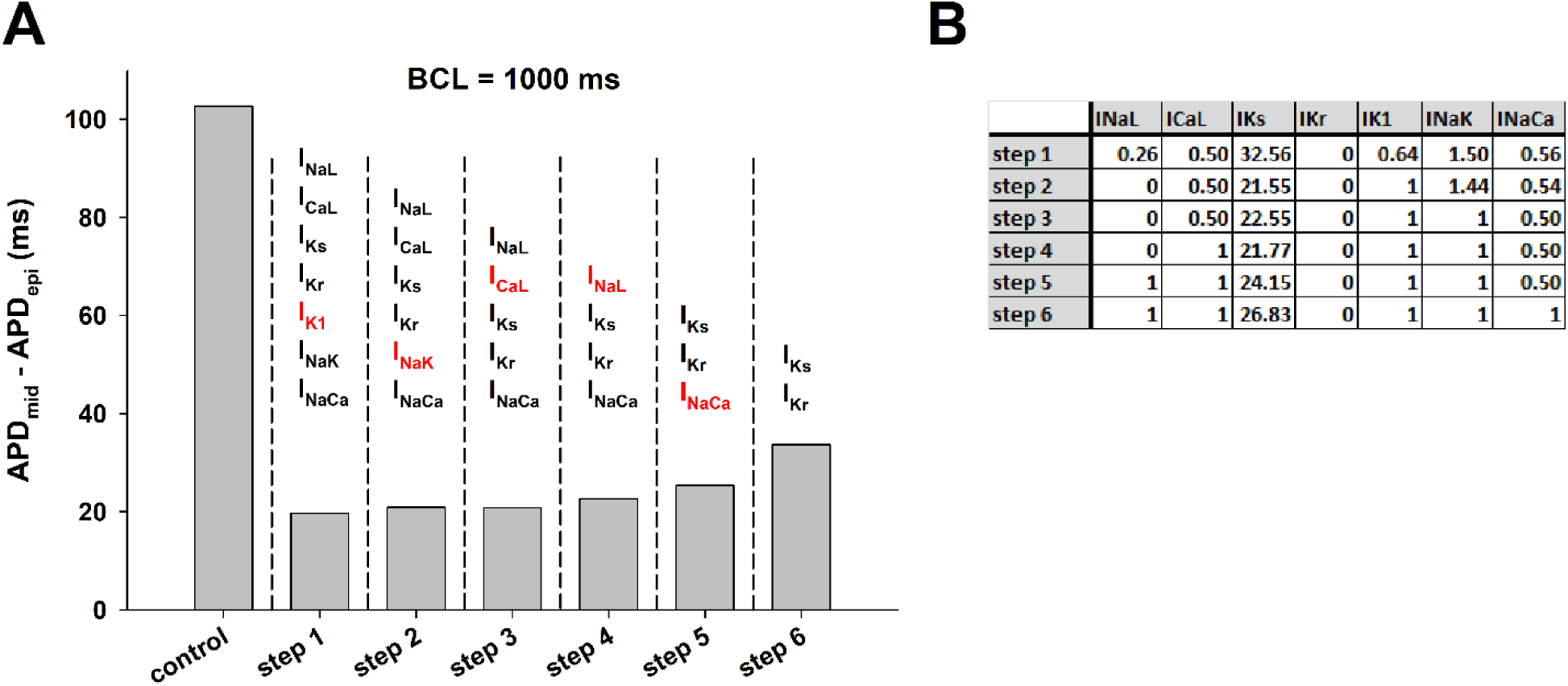
Backward feature elimination procedure applied to the intervention in Figure 8 that reduces dispersion of repolarization when BCL = 1000 ms. **A:** APD dispersion (i.e. the difference between APD in epicardial and mid-myocardial cells) during control and after each step in the procedure. The ion currents that were subject to block are shown on top of each vertical bar. The ion current in red indicates the current eliminated after that step in the procedure because it contributes less to reducing APD dispersion. **B:** Contribution of each ion current at a specific step. See text for detailed description.

Enhancing I_Ks_ and blocking I_Kr_ causes a strong reduction in transmural APD dispersion as a result of the large shortening in mid-myocardial cell APD and a more modest shortening in epicardial cell APD (Figure 9A; Figure 10A and 10B top). To understand the mechanism of the differential APD shortening, we compared the effect on the action potential of an intervention that activates I_Ks_ and blocks I_Kr_ to the control action potential in both types of cells with BCL = 1000 ms (Figure 10). In both types of cells activating I_Ks_ (26.83x the control value) and 100% block of I_Kr_ causes a decrease in the duration of phase 2 (from 160 ms to 132 ms in epicardial cells and from 254 ms to 159 ms in mid-myocardial cells) and an increase in the duration of phase 3 (from 36 ms to 61 ms in epicardial cells and from 47 ms to 68 ms in mid-myocardial cells) of the action potential (Figure 10). The increase in the duration of phase 3 caused by the intervention is about the same in both cells (25 ms in epicardial and 21 ms in mid-myocardial cells). In contrast, the decrease in the duration of phase 2 is much larger in mid-myocardial than in epicardial cells (28 ms in epicardial vs. 95 ms in mid-myocardial cells), which suggests that ion currents during phase 2 are responsible for the differential APD reduction caused by the modulation of the delayed rectifier currents. The difference between the average I_tot_ during control and I_K_ (= I_Kr_ + I_Ks_) modulation in phase 2 is larger in mid-myocardial (control: 0.31 pA/pF/ms; intervention: 0.49 pA/pF/ms; slopes of black and red dotted lines in Figure 10B, top) than in epicardial cells (control: 0.44 pA/pF/ms; intervention: 0.53 pA/pF/ms; slopes of black and red dotted lines in Figure 10A, top). The total delayed rectifier currents are about the same in both cell types during the intervention (I_Ks_ + I_Kr_, red lines in Figure 10A and 10B), but they are considerably smaller in mid-myocardial cells than in epicardial cells during control (I_Ks_ + I_Kr_, black lines in Figure 10A and 10B). As a result, the difference in repolarizing currents between control and intervention is larger in mid-myocardial than in epicardial cells. The larger difference between control and intervention in I_CaL_ (Figure 10A and 10B, bottom) in mid-myocardial cells is not sufficient to compensate for the larger difference in delayed rectifier currents, which explains the larger difference in average total ionic current between control and intervention in mid-myocardial cells, and hence the larger reduction of APD in mid-myocardial cells.

**Figure 10.**
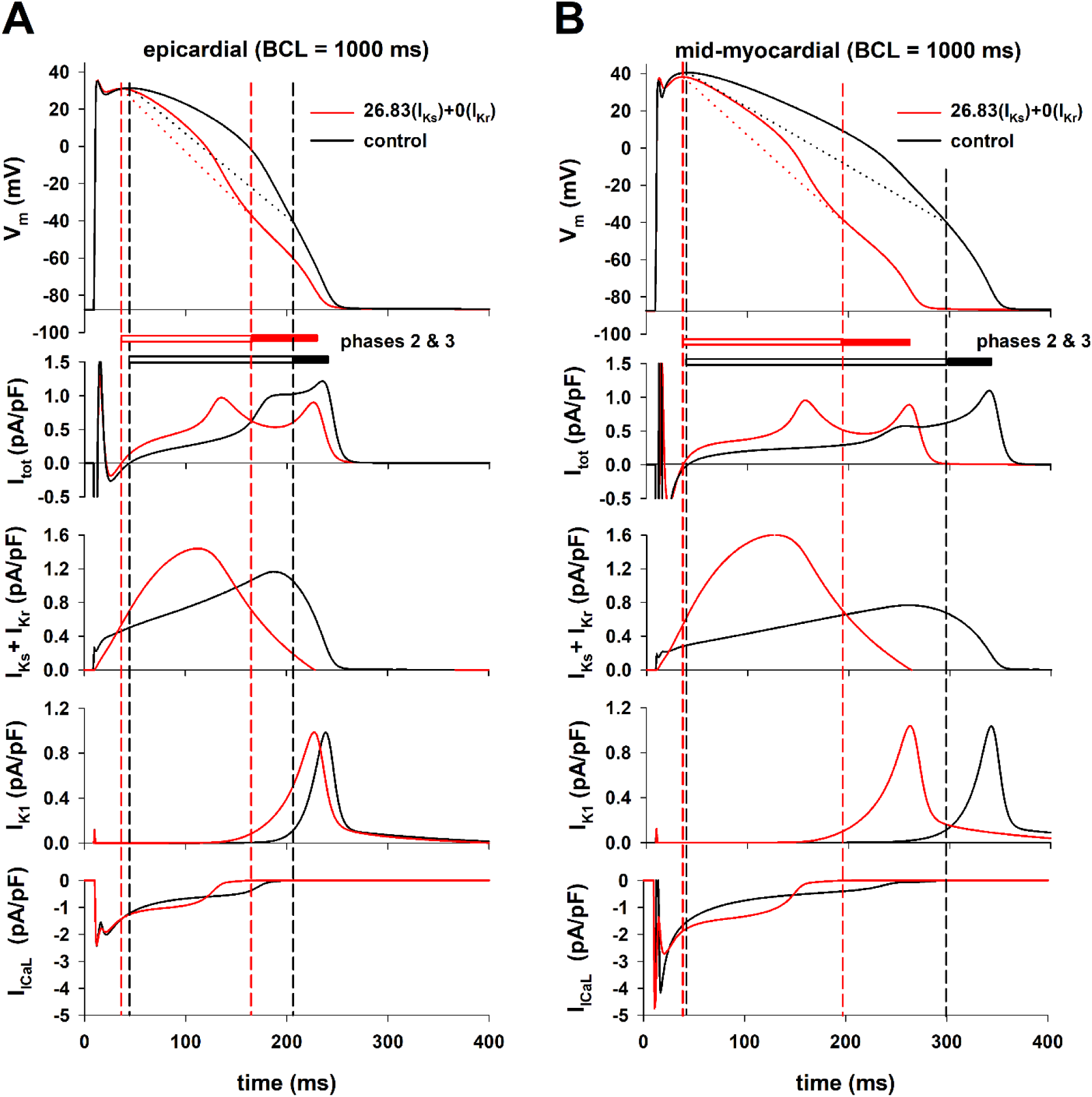
Comparison between the action potential and corresponding ion currents during control (black lines) and an intervention that activates I_Ks_ (26.83 x the control value) and blocks I_Kr_ 100% (red lines) in epicardial (panel A) and mid-myocardial cells (panel B) when BCL = 1000 ms. Black vertical dashed lines indicate the beginning and the end of phase 2 repolarization during control, and the red vertical dashed lines indicate the beginning and the end of phase 2 repolarization during the intervention. See text for detailed description.

### Prolongation of APD without an increase in dispersion of repolarization

In both Figure 4 and Figure 6 the optimal combination of ion channel blockers that minimizes transmural APD dispersion includes 100% block of I_NaL_, 50% block of I_CaL_ and 50% block of I_NaCa_. Figure 11A shows the values of APD of epicardial (black circles) and mid-myocardial cells (black triangles), for different levels of block of I_Kr_ (while maintaining 100% block of I_NaL_, 50% block of I_CaL_ and 50% block of I_NaCa_), with BCL = 1000 ms. In all cases transmural APD dispersion is reduced with respect to control (difference between the dashed lines in Figure 11A), while changing the values of APD of epicardial and mid-myocardial cells, identifying a strategy for prolonging APD and reducing APD dispersion.

**Figure 11.**
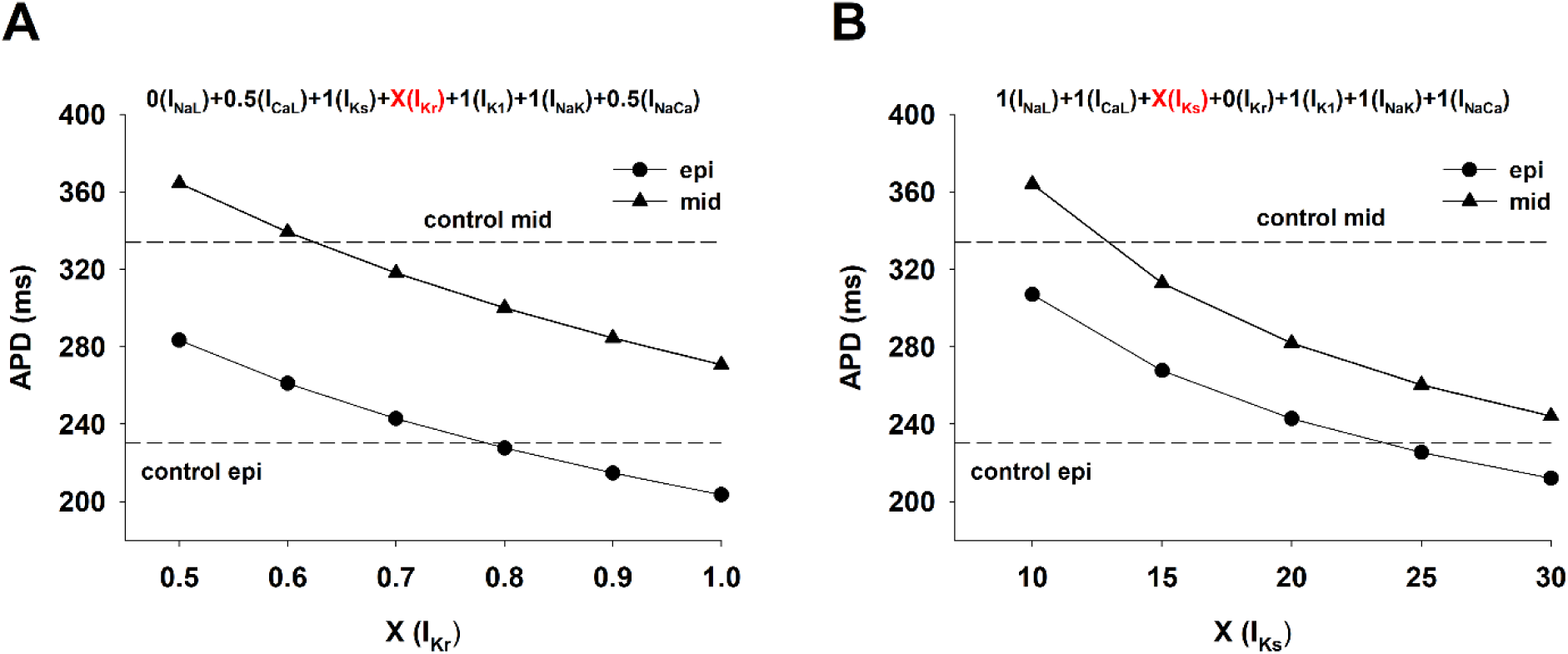
Prolongation of APD with a decrease in dispersion of repolarization. **A:** Interventions that use only ion channel blockers. The plot shows the values of APD for epicardial (black circles) and mid-myocardial (black triangles) cells for different levels of block of I_Kr_ (while maintaining 100% block of I_NaL_, 50% block of I_CaL_ and 50% block of I_NaCa_), with BCL = 1000 ms. **B:** Interventions that use ion channel blockers and activators. The plot shows the values of APD for epicardial (black circles) and mid-myocardial (black triangles) cells for different levels of activation of I_Ks_ (while maintaining 100% block of I_Kr_), with BCL = 1000 ms. Horizontal dashed lines indicate the values of the control APD of epicardial (epi) and mid-myocardial (mid) cells. See text for detailed description.

We have shown earlier that 100% block of I_Kr_ and enhancement of I_Ks_ reduces transmural APD dispersion (Figures 9 and 10). Figure 11B shows the values of APD of epicardial (black circles) and mid-myocardial cells (black triangles), for different levels of enhancement of I_Ks_ (while maintaining 100% block of I_Kr_), with BCL = 1000 ms. In all cases APD dispersion is reduced with respect to control (difference between the dashed lines in Figure 11B), while changing the values of APD of epicardial and mid-myocardial cells, identifying a second strategy for prolonging APD and reducing APD dispersion.

## DISCUSSION

Class III antiarrhythmic drugs primarily act by blocking potassium channels, leading to APD prolongation to prevent reentrant arrhythmias (Peters et al 2000). However, as a result of the heterogeneity in cell types across the myocardial wall, agents that increase APD by selectively blocking repolarizing potassium currents like I_Kr_, may increase APD dispersion, which has been shown to be pro-arrhythmic (Kuo et al 1985; Surawicz 1989; Antzelevitch 2005; Antzelevitch 2008; Avula et al 2019). In this report, using computer models of the action potential of human epicardial and mid-myocardial myocytes, we have identified two strategies to prolong APD while reducing transmural APD dispersion. The first strategy involves blocking several depolarizing and repolarizing ionic currents. Block of I_NaL_, I_CaL_, I_Kr_ and I_NaCa_ can reduce the physiological APD dispersion by about 20% while increasing APD in epicardial and mid-myocardial cells (Figure 11A). The second strategy involves the use of a combination of ion channel blockers and activators which results in a stronger reduction in transmural APD dispersion than using only ion channel blockers. Enhancing I_Ks_ and 100% block of I_Kr_ can reduce the physiological APD dispersion by about 70%, while increasing APD of epicardial and mid-myocardial cells (Figure 11B).

Most of the evidence in the literature suggests that selective I_Kr_ block increases transmural APD dispersion because I_Kr_ blockers preferentially prolong the action potential of mid-myocardial cells (Lukas 1997; Antzelevitch 2008). This increase in dispersion may provide a substrate for the initiation and maintenance of reentrant arrhythmias like TdP (Antzelevitch 2005). Examples include class III selective I_Kr_ blockers like dofetilide and sotalol (Lukas 1997; Antzelevitch 2008). The results of our computer simulations are consistent with those experimental and clinical findings: a 25% block of I_Kr_ prolongs preferentially the APD of mid-myocardial cells, increasing APD dispersion by 8-9% (Figure 1). Despite the larger reduction of average I_tot_ in epicardial cells than in mid-myocardial cells with a 25% block of I_Kr_, the increase in duration of phase 2&3 repolarization (and consequently APD) is smaller in epicardial than in mid-myocardial cells (Figure 5B). This is a consequence of the hyperbolic relationship between average I_tot_ current during phase 2&3 repolarization (I_tot,ph2ph3_) and the duration of phase 2&3 repolarization; the hyperbola has a larger slope for action potentials with longer phase 2&3 (mid-myocardial cells), than for action potentials with shorter phase 2&3 (epicardial cells).

While selective I_Kr_ blockers generally increase transmural APD dispersion, drug agents that block multiple channels in addition to I_Kr_ have been shown to decrease (or at least not to increase) transmural dispersion. For example, ranolazine, which blocks I_NaL_ and I_CaL_ in addition to I_Kr_, has been shown to decrease transmural dispersion (Antzelevitch et al 2004). Also, the reduction in I_NaL_ leads to a decrease in intracellular sodium levels which in turn reduces the reverse mode of I_NaCa_ (Belardinelli et al 2006). Similarly, amiodarone, which is a multichannel acting agent that blocks potassium (I_Kr_, I_Ks_, and possibly I_K1_), sodium (I_Na_ and I_NaL_), calcium (I_CaL_) channels (Sicouri et al 1997; Drouin et al 1998; Vasallo and Trohman 2007; Arpadffy et al 2020; Gelman et al 2024) as well as the sodium/calcium exchanger (Watanabe and Kimura 2000), has also been shown to reduce dispersion of repolarization. In that context, our findings show that optimal multichannel block of I_NaL_, I_CaL_, I_Kr_ and I_NaCa_ can reduce APD dispersion by about 20% (Figures 4, 6, 7, 11A) and are consistent with available experimental and clinical evidence. Figure 11A shows that for an intervention that only blocks depolarizing currents (I_NaL_, I_CaL_ and I_NaCa_, while keeping I_Kr_ at the same value as control (abscissa = 1.0 in Figure 11A), the reduction of APD is more pronounced for mid-myocardial cells (from 334 ms to 271 ms) than for epicardial cells (from 231 ms to 204 ms) thus reducing APD dispersion (from 103 ms to 67 ms). Therefore, while block of repolarizing current I_Kr_ prolongs preferentially APD in mid-myocardial cells (Figure 1), block of depolarizing currents (100% block of I_NaL_, 50% block of I_CaL_ and 50% block of I_NaCa_) shortens preferentially APD in mid-myocardial cells (Figure 11A). All in all, the contrasting and balancing effect of blocking depolarizing and repolarizing ion currents in epicardial and mid-myocardial cells makes it possible to design an optimal strategy to decrease APD dispersion using multichannel blockers. This mechanism may be at work in drug agents like ranolazine and amiodarone that block several depolarizing and repolarizing ion channels and reduce transmural APD dispersion.

The results in Figure 11B show that 100% block of I_Kr_ and enhancement of I_Ks_ can increase APD of epicardial cells while reducing transmural APD dispersion. I_Ks_ activators have been developed to prevent excessive APD prolongation that may occur in patients suffering from LQT syndrome, cardiac hypertrophy, or cardiac failure (Tamargo et al., 2004; Xu et al., 2002; Xu et al., 2015, Bohannon et al 2020). In this report, we have found that the use of I_Ks_ activators in combination with I_Kr_ block decreases transmural APD dispersion. Transmural APD dispersion between epicardial and mid-myocardial cells is mainly the result of the larger I_Kr_ in epicardial cells (Figure 2), which results in a stronger total repolarizing current. The contribution of I_Ks_ to repolarization in both epicardial and mid-myocardial cells is much smaller than that of I_Kr_ (Figure 2). Interestingly, despite its smaller channel density, during the action potential, I_Ks_ is larger for mid-myocardial cells than for epicardial cells (Figure 2). This is a consequence of the morphology of the action potential and the dynamics of activation of I_Ks_. I_Ks_ activates more slowly and at more positive transmembrane potentials than I_Kr_ (Liu and Antzelevitch 1995), and since mid-myocardial cells depolarize to more positive potentials and stay depolarized longer at positive potentials than epicardial cells, I_Ks_ is larger in mid-myocardial than in epicardial cells (Figure 2). Therefore, it is expected that shifting the responsibility of repolarization from I_Kr_ to I_Ks_ would result in a decrease of APD dispersion. Our results are consistent with a computational study by Christophe (2015) who observed that enhancement of I_Ks_ activity results in a decrease in transmural APD dispersion. However, there is experimental evidence suggesting that enhancing I_Ks_ can lead to an increased transmural APD dispersion. Mutations in KCNQ1, which encodes for KvLQT1 a component of I_Ks_ can lead to Short QT Syndrome (SQTS) due to a gain of function in I_Ks_. This gain of function can cause a heterogeneous abbreviation of APD and refractoriness, and an increased inducibility of ventricular fibrillation as a result of an increased APD dispersion (Milberg et al 2007). Beta-adrenergic stimulation can significantly augment I_Ks_ and its contribution to human left ventricular repolarization (Kang et al 2017). They further showed that beta-adrenergic stimulation in combination with I_Kr_ channel blocker E-4031 resulted in significant transmural dispersion of repolarization (Kang et al 2017), which is not consistent with our numerical results.

In conclusion, there is agreement between published experimental and clinical results and the numerical simulations presented in this report indicating that interventions that block several depolarizing and repolarizing ion channels important during phase 2&3 of the action potential can reduce transmural APD dispersion. The numerical simulations in this report provide insight into the mechanisms of that reduction, which ultimately stems from the larger effect on APD of changes in total average ion current (caused by block of either depolarizing or repolarizing ion channels) in mid-myocardial than in epicardial cells. This differential effect is a consequence of the hyperbolic relationship between average total ion current and the duration of phase 2&3 repolarization and can be used to reduce APD dispersion while increasing APD. Conversely, there is no direct experimental or clinical evidence indicating that the second strategy proposed in this report, I_Kr_ blockade in combination with I_Ks_ activation, reduces transmural APD dispersion. The two strategies presented here rely on block of I_Kr_, but there is strong experimental and clinical evidence showing that selective block I_Kr_ leads to an increase in APD dispersion. Therefore, I_Kr_ block should be accompanied by interventions in other ion channels, either by blocking depolarizing currents (first strategy) or activating other repolarizing currents (second strategy).

### Limitations

Computer models of the action potential integrate, often conflicting, experimental data obtained under different conditions from different preparations. As a consequence, conclusions derived from numerical simulations of the cardiac action potential should be interpreted with caution as predictions that need to be tested experimentally. In this report we simulate the effect of anti-arrhythmic agents on ion channels by modulating the maximum channel conductance (see Methods). However, pharmacological agents have binding and unbinding kinetics which may change at different stimulation rates, adding another layer of complexity to their modulation of the channel which was not considered in this report.

## Ethical Approval

This study does not require ethical approval.

